# UBIQUITOUS FUNCTIONAL SYNERGY PARTIALLY EXPLAINS WHY MOST TRANSCRIPTION FACTOR BINDING IS NON-FUNCTIONAL

**DOI:** 10.64898/2026.01.19.700460

**Authors:** Chase Mateusiak, Eric Jia, Jessica N. Plaggenberg, Zolboo Erdenbataar, Yuchen Wang, Christian Shively, Guochen Liao, Robi D. Mitra, Michael R. Brent

## Abstract

Most genes in whose promotor a transcription factor (TF) binds do not change in expression when the concentration of the TF is perturbed. No existing model can predict which bound promotors will respond and which will not. We hypothesized that a gene’s response to perturbation of a TF bound in its promotor can depend on which other TFs are bound there, a phenomenon we call functional synergy. This is distinct from cooperative binding, which is already accounted for in the binding location data. To investigate functional synergy, we created a comprehensive dataset on TF binding locations in yeast using a method that is orthogonal to chromatin immunoprecipitation. We then used mathematical modeling to identify high-confidence instances of functional synergy. We found that such synergies are surprisingly common. Responses to perturbations of 44 different TFs were modified by the presence of other TFs. 48 TFs served as modifiers, but some modified responses to many TFs. We conclude that (1) measuring the binding locations of a single TF will not, in general, reveal which genes the TF regulates, and (2) traditional networks linking TFs to their targets must be made substantially more expressive, allowing some TFs to modify the effects of others.

## INTRODUCTION

A surprising, but highly robust and often-replicated observation is that only a small fraction of the genes whose regulatory DNA is strongly bound by a transcription factor (TF) respond to perturbations of that TF with a measurable change in transcript abundance. This has been shown in multiple datasets on yeast^8–12^ and human cells.^8,13^ Currently, there are no models that can explain which binding events affect transcription and which do not. This paper begins with the question, “Why do some genes whose promoter are strongly bound by a TF respond to perturbations of that TF, while others do not?”

One hypothesis is that non-functional binding sites represent technical artefacts of chromatin immunoprecipitation (ChIP), the predominant methodology for measuring binding locations. Indeed, ChIP has many well-documented limitations. For example, ChIP experiments targeting a protein that is not expected to bind DNA, such as green fluorescent protein, show thousands of ChIP peaks in the same regions where peaks for many TFs cluster – regions with high GC content, open chromatin, and highly transcribed genes^18–20^. Similar peaks are found in ChIP experiments targeting proteins that are not present in the sample^22,23^. These may arise in part because DNA fragmentation is biased toward breaking DNA in open chromatin and near the 5’ ends of genes, especially highly expressed genes^25^. These regions appear on shorter segments of post-sonication DNA, on average. Preparation of DNA for hybridization or sequencing involves selecting for short fragments, which enriches for regions of open chromatin near highly expressed genes,^25^ precisely where non-specific ChIP signals are seen.

To investigate the possibility that non-functional binding sites are artefacts of ChIP, we produced a comprehensive dataset of TF binding locations in yeast, described here for the first time, using Transposon Calling Cards (TCC).^30–33^ TCC is very different from ChIP in that it involves no physical DNA fragmentation, no affinity purification, and no protein handling. Furthermore, binding locations are recorded in living cells in which TF-DNA binding is approximately at equilibrium. To carry out the TCC assay on a yeast TF, the Sir4 domain, which attracts the Ty5 transposon, is fused to the TF (Fig. 1A). Some fraction of the time when the TF is bound to DNA, it inserts a transposon nearby. These transposons are recovered, along with flanking DNA, and sequenced to reveal the locations of the insertions. We report below that, while our CC data do show better agreement with perturbation responses than older ChIP data, it is still the case that most binding is non-functional, even in a genome as compact and simple as the yeast genome. Thus, our TCC results rule out the technical artifacts associated with ChIP as a primary explanation for why so few of the genes whose promoters are strongly bound by a TF respond to perturbation of that TF.

**Figure 1.**
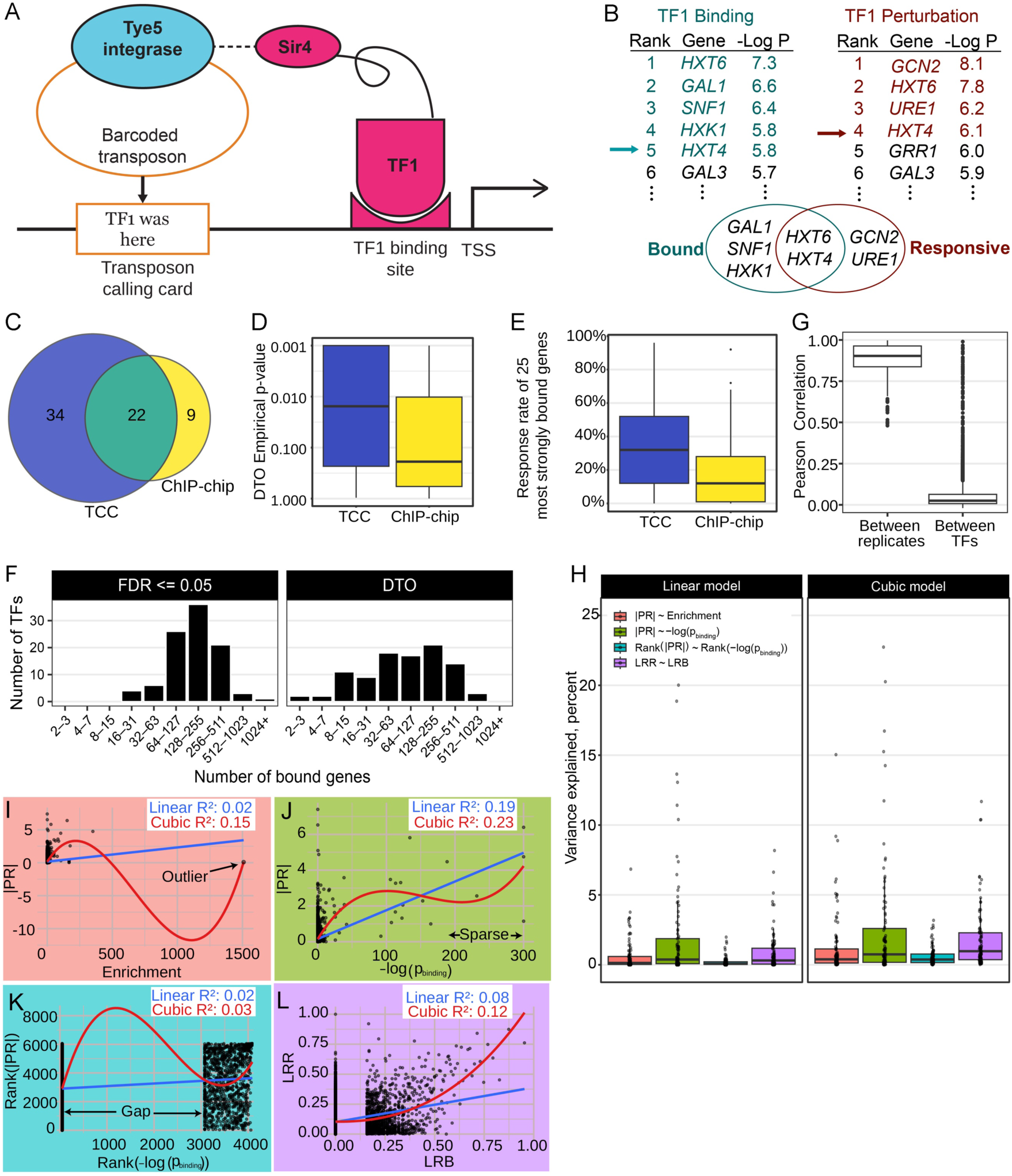
(A) Illustration of the Transposon Calling Cards method. (B) Illustration of the Dual Threshold Optimization (DTO) algorithm. Arrows indicate thresholds on binding signal rank (turquoise) and perturbation response rank (maroon). Thresholds are chosen to maximize the hypergeometric P-value of the overlap between bound and responsive genes. A permutation-based distribution is used to test for statistical significance of the overlap. (C) Venn diagram of TFs with significant bind-perturbation overlap, determined by DTO. TCC, transposon calling cards. Only TFs present in both datasets are considered. (D) Distributions of DTO empirical P-values for TCC and ChIP-chip data. (E) Percent of genes that respond to perturbation of a TF among the 25 genes whose promoters are most strongly bound by the TF (see Methods). (F) Distributions of number of bound targets per TF, as determined by FDR<= 0.05 or by DTO empirical P-value <= 0.01. (G) Correlations between replicates of the same TF in our calling cards data (left) and between pairs of distinct TFs (right). Each point represents a pair of TFs. (H) Distribution of variance explained (R^2^) when modeling various transformations of the data using linear models (left) or cubic models (right). Each point is a model for responses to perturbation of one TF. (I-L) Responses of genes to perturbation of Ytf1 (vertical axis) and strength the binding signal in the genes’ promotors (horizontal axis). In each panel, data are transformed differently (see main text. Blue: linear model fit. Red: model fit. Each point represents one gene.

We then used our new data to investigate the hypothesis that a gene’s response to perturbation of a TF that is bound in its promoter depends on which other TFs are bound there. We refer to such dependencies as functional synergies, to distinguish them from cooperative binding. Cooperative binding is important for explaining <u>why TFs bind</u> where they do, but functional synergy is an attempt to explain the effect on transcription, or lack therefore, when a TF <u>is</u> bound. We find that functional synergies between bound TFs are common and involve many different TFs, although certain TFs appear especially prone to modify the effects of others. Specifically, computational models for predicting the response of a gene to perturbation of a TF that is bound in its promotor are much more accurate when they use information about which other TFs are bound in its promotor. To the best of our knowledge, no previous research has attempted to predict the response to perturbation of one TF by using the binding locations of other TFs.

## RESULTS

### A new, genome-wide dataset on TF binding locations

We carried out multiple replicates of the calling cards experiment on 230 different yeast TFs (see Methods). Briefly, strains expressing a TF tagged with Sir4 and a retrotransposon donor plasmid with expression driven by the *GAL1* promoter were incubated on agarose with synthetic complete medium and galactose as the sole carbon source (Fig. 1A). After 4 days, cells were recovered from agarose plates and DNA was extracted and cut with three restriction enzymes. The resulting fragments were self-ligated and subjected to inverse PCR, using primers that are complementary to the transposon, followed by Illumina sequencing. Sequencing reads from each replicate for a given TF were analyzed using our custom pipeline (https://nf-co.re/callingcards),^33^ which counts insertions in the promoter regions of genes and compares the counts in each promotor to those of insertions in the same promoter in cells lacking Sir4. For each TF and each gene’s promoter, our pipeline outputs a P-value for the null hypothesis that the probability of a transposon being inserted in the strain with the Sir4-tagged TF is no greater than in the sir4-deleted strain.

Next, we set out to determine whether the low response rate among genes whose promoters are bound by a TF could be explained by artefacts of the ChIP methodology. To do this, we compared both ChIP data^36^ and our CC data to changes in gene expression after overexpression of TFs using the ZEV system.^37^ Such comparisons often made by setting a P-value threshold on binding to identify a set of bound genes and on differential expression to identify a set of responsive genes. However, the results of such comparisons are highly sensitive to the P-value thresholds and statistical methods chosen. Furthermore, they fail to take advantage of information that each data type provides about which results in the other data type are biologically meaningful. For example, a binding signal of uncertain significance is more likely to reflect a biologically meaningful binding if it falls in the promoter of a gene that responds to perturbation of the same TF. To take advantage of this, we developed Dual Threshold Optimization (DTO), which considers all possible pairs of thresholds on the two data types and selects the pair that maximizes the significance of the overlap between the bound and responsive sets (Fig. 1B).^8^ The DTO software^38^ then calculates an empirical, permutation-based p-value for the overlap, which we consider to be significant when 𝑃 ≤ 0.01.

To make a fair comparison between calling cards and ChIP, we focused on the 118 TFs that are present in both binding datasets and in the TF overexpression data. Of the 118 TFs, 31 had a DTO empirical p-value ≤ 0.01 in the ChIP data, whereas 56 did in the TCC data (Figs. 1C). In fact, the entire P-value distribution was shifted toward greater significance (Fig. 1D) in the TCC data. Thus, TCC tends to agree with the perturbation response data better than the ChIP data does. However, it remains the case that TCC data on 66 of these TFs (56%) does not show a significant overlap between bound and responsive genes. Among TFs that do show a significant overlap, the median fraction of bound genes that are responsive is 20%, based on the DTO thresholds (not shown). Even for the 25 most strongly bound targets of each TF, using the authors’ threshold for responsiveness, the median response rate is only 32% (Fig. 1E, Methods). Thus, we still need to explain why some genes whose promoters are bound by a TF respond to perturbation of that TF while others do not.

Before analyzing the TCC dataset further, we wanted to eliminate replicates that showed little to no relationship between binding locations and perturbation responses. To this end, we ran DTO comparing calling cards data to both the TF overexpression data (TFOE) and to a different dataset on responses to TF knockout (TFKO).^39^ We used a TCC replicate if DTO analysis produced an empirical P < 0.01 compared to either perturbation dataset; other replicates were discarded, leaving data on 122 different TFs (some of which bind DNA as part of a complex rather than directly), with a median of 2 replicates per TF. In subsequent analyses, reads from all usable replicates for a TF were combined and run through the analysis pipeline again. We then calculated binding P-values for each TF-gene pair (see Methods) and found that the median number of significantly bound targets per TF was 146 (FDR<=0.05). Figure 1F shows the distribution of number of targets per TF based on FDR<0.05 (left) and based on the DTO thresholds after comparison to the TFOE data (right). In both cases, the mode was between 128 and 255 bound targets, with the DTO analysis showing a broader, flatter distribution than the FDR analysis, which is based on TCC data without considering perturbation responses.

We also calculated the correlation of –log *P* between replicate pairs for the same TF, where *P* is the *P*-value for the TF binding in a gene’s promoter. The median correlation across all replicate pairs was 0.90. We then combined all replicates for each TF and calculated the correlations between all pairs of TFs (Fig. 1G), yielding a median of 0.025. For the 24 pairs of TFs with the highest correlation (> 0.81), there were strong biological reasons to expect that they would have similar binding profiles (Table 1); the fact that they do have similar binding profiles gives us confidence in the quality of our data.

**Table 1:**
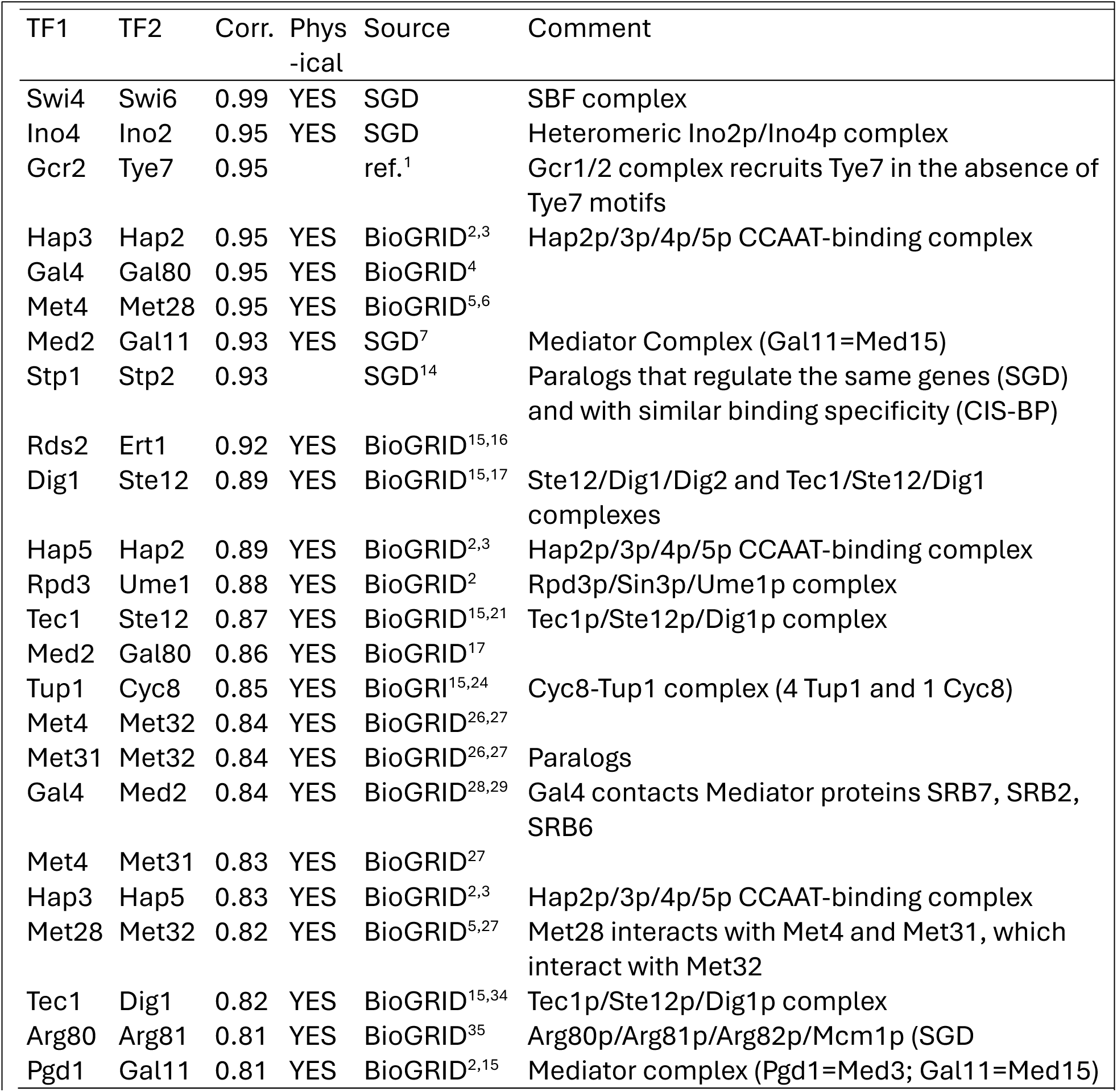
The TFs with most correlated binding profiles and why their profiles are expected to be similar.

### Data transformations and univariate regression models

Next, we set out to model the relationship between binding and perturbation response. To better reflect the underlying biochemistry, we treated both binding strength and perturbation response strength as continuous variables and used a regression approach. We built a separate model for each TF because: (1) TFs differ in their activation or repression strengths and their activity levels in the experimental conditions tested, (2) we hypothesized that different TFs would have different synergy partners, and (3) separate models give us the best chance of finding meaningful relationships between binding strength and response strength.

As a baseline for comparison, we first fit a univariate linear model to predict the absolute value of each gene’s log fold change (LFC) in response to perturbation of a TF by using the binding signal for the TF in the promoter of the gene (see Methods). For this and subsequent analyses, we built separate models for the responses to perturbations of the 97 TFs for which we had both QC-passing TCC data and TFOE data. Initially, the *binding signal* we used for modeling was enrichment: the ratio of observed transpositions into a promoter in the TF-Sir4 fusion strain compared to the number of transpositions in the same promoter in a control strain lacking Sir4. Data on 52 of the 97 TFs showed a statistically significant linear relationship between binding locations and perturbation responses with *P* < 0.01 (F-test). The distribution of variance explained for the models across all TFs (Fig. 1H, left, red box) shows that the linear models explain only a tiny fraction of variance in the data (median = 0.13%). We then fit cubic polynomials, in case only the most strongly bound genes are more responsive than the rest. This improved the distribution of variance explained across TFs (Fig. 1H, right, red box). However, plotting the data showed that, in many cases, the fits were unduly influenced by outlier genes with huge binding enrichment (Fig. 1I, arrow). Furthermore, very high enrichments can be achieved with relatively few transpositions, if there are no transpositions into the promoter in the control strain.

To combat both outliers and low counts, we next tried using the -Log *P*-value as the binding signal. This improved the variance explained by the linear and cubic models (Fig. 1H, green) and the distribution of binding signals, but the data points remained sparse. For example, in Figure 1J, the top third of the binding signal distribution, between 200 and 300, contains only four datapoints. Next, we tried using the rank of the -Log *P*, with the least significant *P*-value receiving rank 1 and the most significant receiving the largest rank. There are many ties, especially at *P*=1.0. This led a big gap between genes with a P-value of 1.0 and the rest and relatively little separation among, for example, the 300 most strongly bound genes, where differences are less likely to reflect measurement noise (Fig. 1K). The cubic fit shows wild swings because there are no data points to constrain it in the large gap. Finally, the variance explained distribution was even worse than before (Fig. 1H, turquoise).

Next, we broke rank ties among P-value ties using enrichment (see Methods for details). This mainly affects small P-values since there are no transpositions, and therefore no enrichment, for most genes. We then took the logarithm of the ranks (equivalent to an inverse exponential transformation) of both the binding P-value and the absolute log fold change of the response. Finally, we divided the log ranks by the log of the number of genes in the yeast genome so that the final binding and response signals lay in the range between zero and one (Fig. 1L). We refer to these transformed signals as the log rank binding (LRB) and log rank response (LRR). The log rank transformation emphasizes strong signals (small P-values), which are least likely to be dominated by noise, and compresses weak signals, which are more likely to be dominated by noise. It can therefore be viewed as a less extreme version of binarizing the data based on a P-value threshold, which is the transformation typically applied to these data. For example, in Fig. 1K, there are six genes in the top quartile of the LRB range, 51 in the next quartile, and 5,591 in the bottom quartile. This makes sense because we expect that most promoters are not occupied by any given TF at biologically meaningful levels. There is still a gap between the genes with binding *P*=1.0 (LRB=0) and the rest of the genes, but it is now much smaller. This transformation often led to the types of fits we expected, where the predicted response is a monotonically increasing function of binding strength and each portion of the fit line is influenced by many points, not just a few outliers (Fig. 1L). It also increased the variance explained, relative to the untransformed data, reflecting an improved handling of outliers and weak binding signals (Fig. 1H, purple). Supplemental Files S1 and S2 contain the LRB and LRR transformed data, respectively. Supplemental Files S3 and S4 contain the coefficients and residuals from the cubic models of LRR as a function of the LRB of the perturbed TF, respectively.

### Modeling functional synergy

Even after fitting models that predict the perturbation response as a function of the binding signal with cubic polynomials, no model for any TF explains more than 12% of the variance in the response data (R^2^), and the median R^2^ is less than 1% (Fig. 1H, purple). As expected from previous studies, the binding signal from the perturbed TF (pTF) has limited power to predict the perturbation response. This implies that other factors, in addition to the binding of the pTF, influence the response. We hypothesized that the missing factors are other TFs bound at the same promoter and set out to investigate that possibility using mathematical modeling.

To focus on the variance in responses that cannot be explained by the binding of the pTF, we regressed out the effect of pTF binding by extracting the residuals from the cubic fits and building new models to explain those residuals. This protects against other variables appearing to be predictive only because they are correlated with (or otherwise related to) the binding of the pTF. We then built the simplest linear statistical models that could represent functional synergy between bound TFs; namely, a model with interaction terms between the binding signal of the pTF and the binding signals of all other TFs:

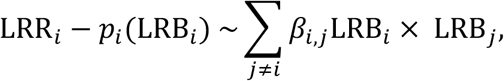

where LRR_𝑖_ is the log rank response of a gene to perturbation of TF_𝑖_, LRB_𝑖_ is the log rank binding signal of TF_𝑖_ in the gene’s promoter, and 𝑝_𝑖_(LRB_𝑖_) is the predicted response from the cubic fit. The left hand side of the equation is thus the residual from the cubic fit. The summation is over the binding signals of all other TFs, and 𝛽_𝑖,j_ is the coefficient of the interaction term for LRB_𝑖_ and LRB_j_. The interaction term itself, LRB_𝑖_ × LRB_j_, is large only when both TF_𝑖_ and TF_j_ are large. At promoters where TF_𝑖_ is not bound (LRB_𝑖_ ≈ 0), no other TF affects the response to perturbation of TF_𝑖_. Thus, this model focuses on the effect of both TF_𝑖_ and TF_j_ being bound in the same promoter.

If we included a term for LRB_j_ alone, that would represent the independent effect of TF_j_ binding, regardless of whether the pTF (TF_𝑖_) is bound in the same promoter. Such terms are appropriate for modeling indirect effects of a perturbation of TF_𝑖_ that are mediated by its effect on the expression of TF_j_, but that is not our goal here. In our model, 𝛽_𝑖,j_ ≠ 0 indicates functional synergy -- the presence of TF_j_ changes the response of promoters where TF_𝑖_ binds to perturbations of TF_𝑖_. If 𝛽_𝑖,j_ > 0, the presence of TF_j_ enhances the response to perturbations of TF_𝑖_; if 𝛽_𝑖,j_ < 0, the presence of TF_j_ inhibits the response to perturbations of TF_𝑖_. Because we are modeling the absolute value of the response to perturbation of TF_𝑖_, this is true regardless of whether TF_𝑖_ activates or represses the gene. When 𝛽_𝑖,j_ is significantly different from zero, we call TF_j_ a modifier TF (mTF) of TF_𝑖_, the perturbed TF (pTF).

### A stringent pipeline for identifying robust functional synergies

We implemented a stringent, three-stage pipeline for identifying the most significant and robust indicators of functional synergy (Fig. 2A). All stages used L1 regularized linear regression (LASSO). LASSO tends to set the coefficient of each feature (interaction term) to 0 unless it has substantial predictive power in cross validation. In Stage 1, we modeled the responses of all genes. In Stage 2, using only mTFs that survived the first stage, we modeled the responses of the 600 genes that are most strongly bound by the pTF, in order to weed out any significant interactions from Stage 1 that were driven primarily by weak signals.

**Figure 2.**
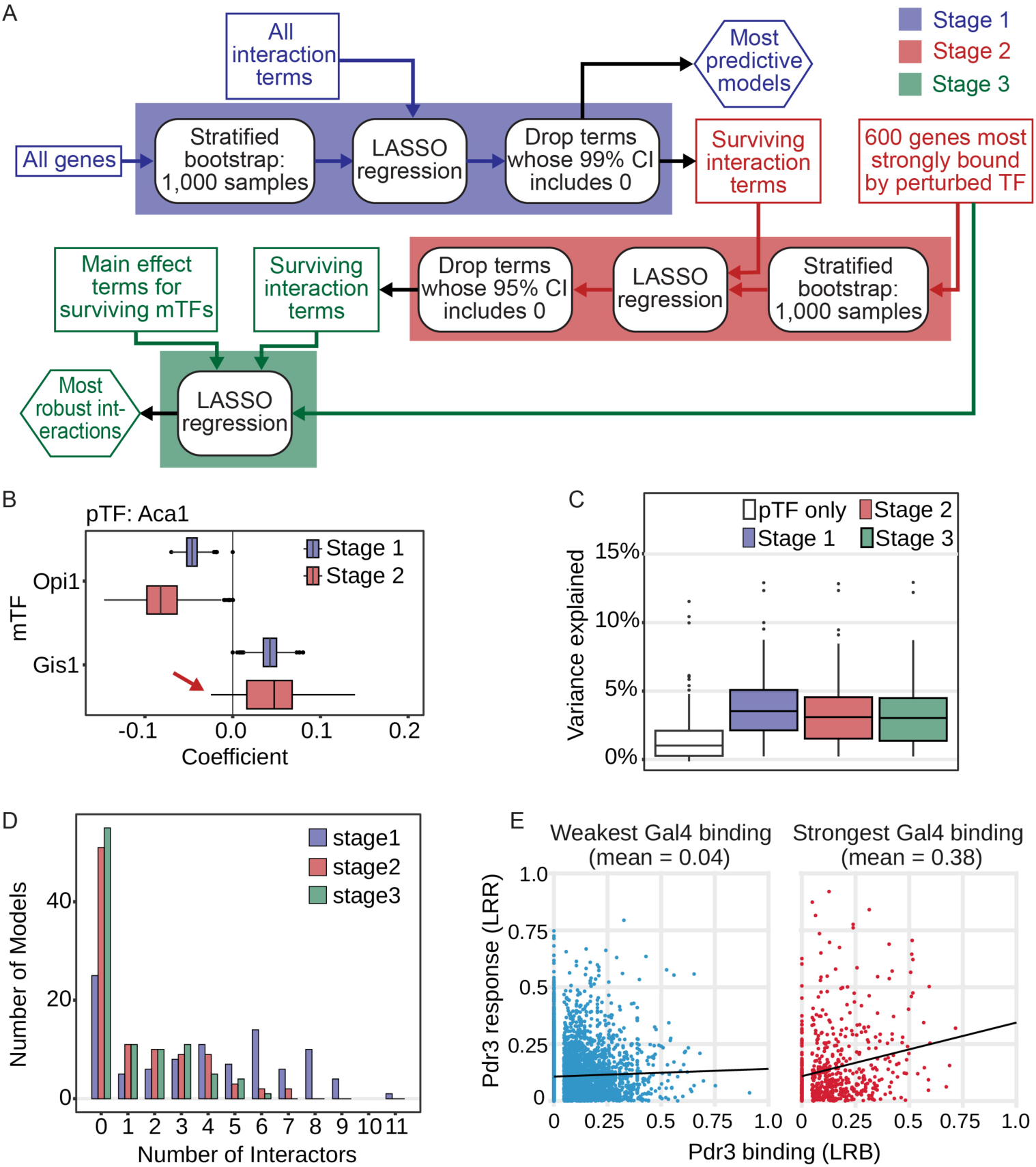
(A) Diagram of the three stages of our pipeline for identifying robust, high-confidence functional synergy. (B) Distribution of coefficients for the interaction terms Aca1*OpI1 and Aca1*Gis1 as predictors of the response to perturbation of Aca1. Each point is a model fit to one of 1,000 bootstrap samples. Both terms survive Stage 1, but Aca1*Gis1 because more than 5% of bootstrap samples have 0 or negative coefficients. (C) Distributions of variance explained, estimated by cross-validation, when models including interaction terms that survive stage 1, or Stages 1 and 2, or Stages 1, 2, and 3, are fit. As expected, variance explained is slightly reduced when terms that do not survive Stage 2 or Stage 3 are removed. Each point is one model. (D) Number of surviving interaction terms after Stages 1, 2, and 3. (E) Visualization of the significant effect of Gal4 binding as a modifier of the response to perturbation of Pdr3. Left: At promoters where Gal 4 is bound weakly or not at all, the strength of Pdr3 binding is not a good predictor of the response to Pdr3 perturbation. Right: At promoters where Gal 4 is more strongly bound, stronger Pdr3 binding predicts stronger response to perturbation of Pdr3.

In Stages 1 and 2, we created 1,000 bootstrap datasets by sampling data points (genes) at random with replacement to obtain a new sample with the same number of genes. Next, we ran LASSO on each bootstrap sample. Using the distribution of coefficients for each term across all bootstrap samples, we created confidence intervals (Fig. 2B). In Stages 1 and 2, we iteratively eliminated terms that were retained in < 50% of bootstrap samples and fit a new LASSO model with the remaining terms to the bootstrap samples, until no more terms were eliminated. In Stage 1, we then applied a 99% one-sided confidence interval and filtered out terms whose intervals included zero. In Stage 2 we applied a 95% one-sided confidence interval, since we only used 10% of the data and our goal was to provide an extra check on terms that have already been shown to be significant in the full gene set. The models resulting from Stages 1 and 2 can be found in Supplemental Files S5 and S6.

In Stage 3, we fit a LASSO model with terms for all interactions that survived Stage 2 and the corresponding independent effects of the mTFs’ binding signals. If LASSO dropped an interaction term in the presence of the corresponding independent effect, we considered that the interaction might have been included because of its correlation to the independent effect term, which could represent an indirect effect mediated by the mTF rather than a functional synergy. Such terms were dropped from the list of high confidence functional synergies. Modifier TFs whose coefficients changed signs (from enhancing to attenuating the response, or vice-versa) were also dropped. Finally, the main effects terms were removed and models consisting of the surviving interaction terms were refit to the data (Supplemental File S7).

By normal statistical criteria, the coefficients of the interaction terms retained after Stage 1 would be considered highly significant, based on the 99% bootstrap confidence intervals. As expected, the models resulting from Stage 1 are the most accurate predictors of the form shown in the equation (Fig. 2C, purple). Stages 2 and 3 eliminated terms for which there might be an explanation other than functional synergy. The resulting models are less accurate predictors of perturbation responses, but the surviving interaction terms represent higher confidence functional synergies. However, interaction models from all stages explained significantly more variance (estimated by cross validation) than the univariate cubic models alone (Fig. 2C).

### Functional synergy is surprisingly common, involving many modifier and modified TFs

The most important finding from this investigation is the surprisingly high frequency with which the presence of an mTF in promotor modifies the response to perturbation of the pTF. After Stage 1, 75% of models included at least one significant modifier term, after Stage 2 47% did, and after Stage 3, 45% did. For pTFs that had at least one mTF after Stage 3, the median number of modifiers was 2.5 (Fig 2D). Table 2 presents the mTFs for each pTF. Figure 2E shows an example of a significant modifier effect. Both plots show the Pdr3 perturbation response as function of Pdr3 binding strength, but for different subsets of genes. On the left are the genes that are bound weakly or not at by Gal4. For these genes, the strength of Pdr3 binding has essentially no effect on the response to Pdr3 perturbation, as shown by the nearly horizontal trend line. On the right are the 600 genes that are most strongly bound by Gal4. For these genes, stronger Pdr3 binding leads to stronger response to perturbation of Pdr3. The difference in slope between the trend lines on the left and the right represents the interaction between Pdr3 binding and Gal4 binding. Figure S1 provides an alternative visualization of the interaction. In total, 48 different TFs modify the responses to perturbations of other TFs. Our second important finding is that some TFs serve as modifiers much more often than the rest, including Msn2, Gal4, Gis1, and Opi1 (Fig. 3A; see Discussion). Interestingly, mTFs that modified three or more pTFs affected most or all of them in the same direction (Fig. 3B). For example, the presence of Msn2 increased responses to perturbations of 11 pTFs while decreasing the responses to only two.

**Fig. 3.**
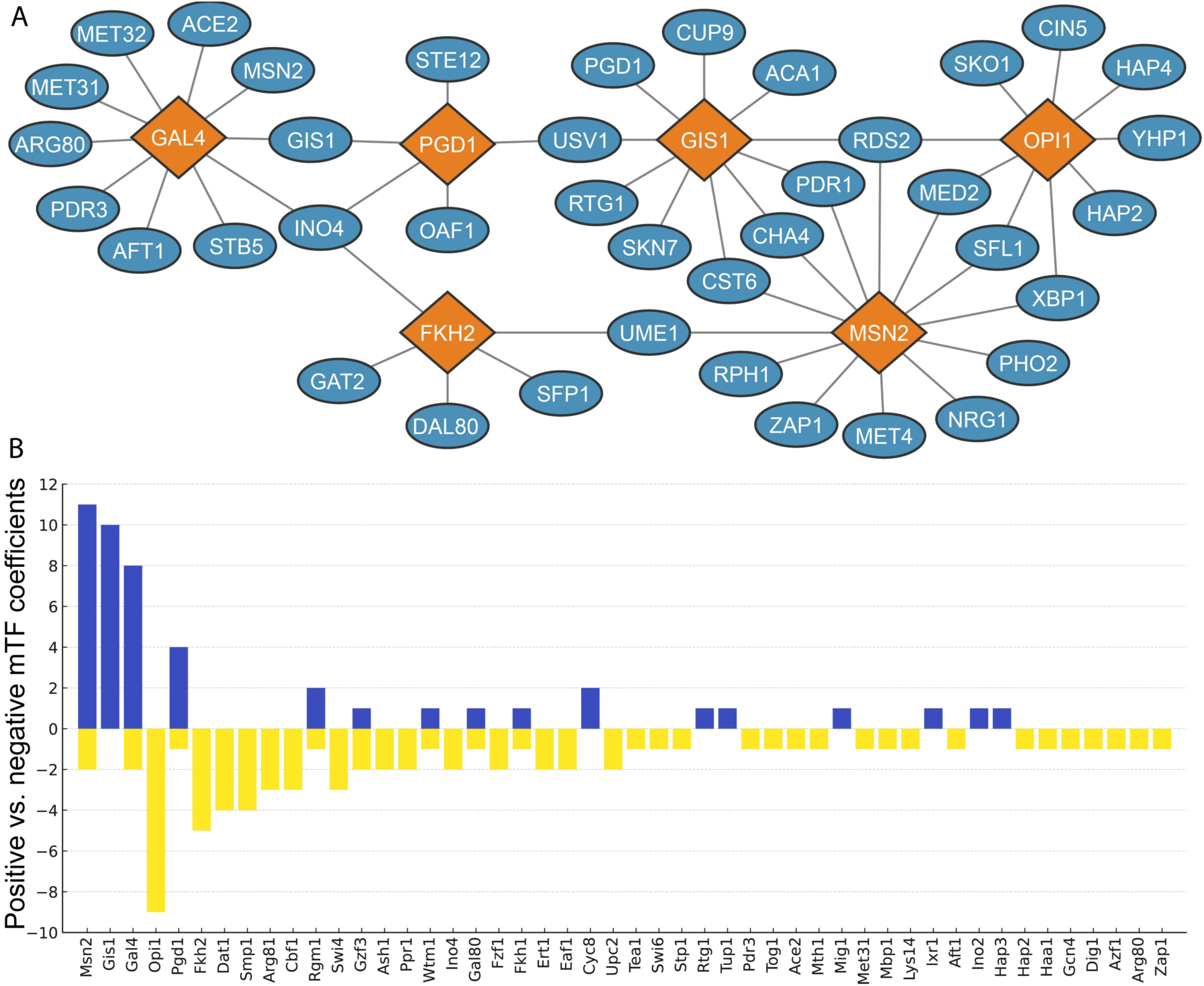
A. Modifier TFs (orange) whose binding locations modify responses to perturbations of the most pTFs (blue). B. For each TF that served as mTF, the number of times its coefficient was positive or negative.

**Table 2.** For each perturbated TF model, modifier TFs that survived all stages of the selection pipeline.

| pTF | mTFs | pTF | mTFs | pTF | mTFs | pTF | mTFs | pTF | mTFs | pTF | mTFs | pTF | mTFs |
| --- | --- | --- | --- | --- | --- | --- | --- | --- | --- | --- | --- | --- | --- |
| **Rds2** | Gis1<br>Gzf3<br>Msn2<br>Opi1<br>Rgm1<br>Swi4<br>Upc2 | **Cha4** | Gis1<br>Lys14<br>Msn2<br>Pdr3<br>Rtg1 | **Dal80** | Cbf1<br>Eaf1<br>Fkh2<br>Gzf3<br>Mth1 | **Gal11** | Aft1<br>Cyc8<br>Gcn4<br>Stp1<br>Zap1 | **Hap4** | Gal80<br>Hap2<br>Hap3<br>Opi1<br>Smp1 | **Pdr1** | Arg81<br>Gis1<br>Gzf3<br>Msn2<br>Upc2 | **Rph1** | Dat1<br>Ert1<br>Msn2<br>Rgm1<br>Tog1 |
| **Cst6** | Arg81<br>Ash1<br>Gis1<br>Msn2 | **Hap2** | Dat1<br>Fkh1<br>Mig1<br>Opi1 | **Ino4** | Fkh2<br>Gal4<br>Pgd1<br>Smp1 | **Met31** | Dig1<br>Gal4<br>Smp1<br>Tup1 | **Sfl1** | Fzf1<br>Msn2<br>Opi1<br>Swi4 | **Usv1** | Azf1<br>Gis1<br>Pgd1<br>Wtm1 | **Xbp1** | Gal80<br>Ino4<br>Msn2<br>Opi1 |
| **Cin5** | Eaf1<br>Ixr1<br>Opi1 | **Cup9** | Ert1<br>Fkh1<br>Gis1 | **Gis1** | Arg80<br>Gal4<br>Pgd1 | **Med2** | Met31<br>Msn2<br>Opi1 | **Oaf1** | Mbp1<br>Pgd1<br>Rgm1 | **Opi1** | Fzf1<br>Ino2<br>Ppr1 | **Pgd1** | Arg81<br>Gis1<br>Swi6 |
| **Skn7** | Gis1<br>Haa1<br>Ppr1 | **Ume1** | Fkh2<br>Ino4<br>Msn2 | **Aft1** | Dat1<br>Gal4 | **Gat2** | Dat1<br>Fkh2 | **Met32** | Cbf1<br>Gal4 | **Met4** | Msn2<br>Smp1 | **Mig1** | Cbf1<br>Cyc8 |
| **Msn2** | Ash1<br>Gal4 | **Sfp1** | Fkh2<br>Tea1 | **Zap1** | Msn2<br>Swi4 | **Aca1** | Gis1 | **Ace2** | Gal4 | **Arg80** | Gal4 | **Nrg1** | Msn2 |
| **Pdr3** | Gal4 | **Pho2** | Msn2 | **Rpn4** | Ace2 | **Rtg1** | Gis1 | **Sko1** | Opi1 | **Stb5** | Gal4 | **Ste12** | Pgd1 |
| **Yhp1** | Opi1 | **Yox1** | Wtm1 |  |  |  |  |  |  |  |  |  |  |

### Converting mathematical models into a new type of TF network map

Once the parameters of our models have been fit to the data, they can take in binding data measured under new conditions and predict the regulatory effects of TFs on target genes in those conditions. These effects can be visualized as networks in which the relationships between TFs and their targets are regulated by other TFs (Fig. 4). These diagrams depict the independent regulatory effect of a TF on a target as a link between the TF and the target, with additional links from modifier TFs into the primary link. Each link can be quantified by its contribution to the target’s predicted response to perturbation of the primary regulator (Fig. 4, edge width). These diagrams improve on traditional TF network maps in two important ways (see Discussion):

1. They can represent complex relationships between TFs and targets, whereas traditional network maps represent only relationships between a single TF and target.
2. The regulatory effects depicted are derived by feeding binding data into a mathematical model, so they are specific to the conditions in which binding locations were assayed.

**Figure 4.**
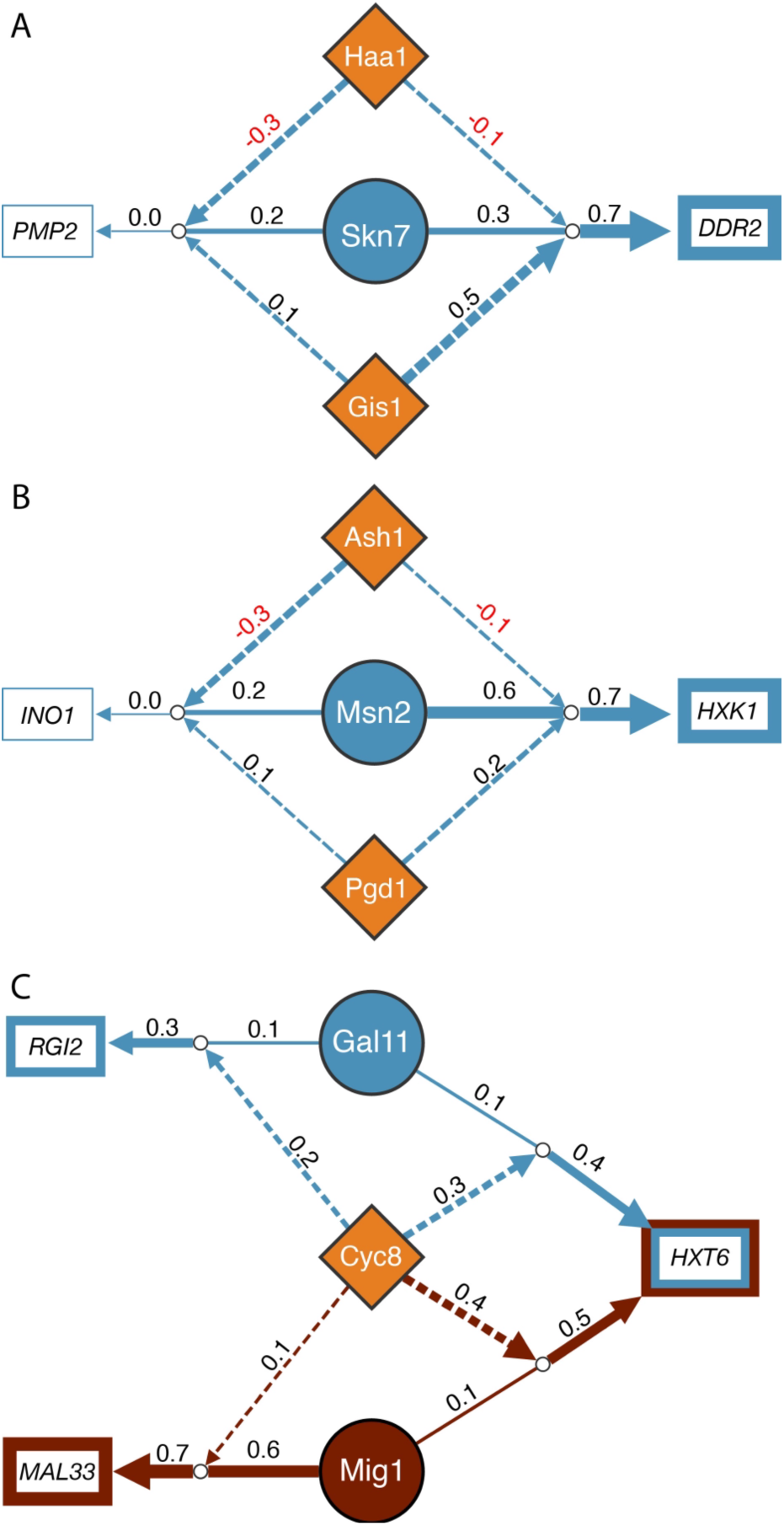
Network diagrams of regulatory effects. Rectangles, target genes. Large circles, perturbed TFs. Diamonds, modifier TFs (mTFs). Line weights and annotations indicate predicted contributions to the total effect of perturbing the pTF of the same color on the target. Dashed lines: contributions of interactions. (A) Perturbation of Skn7 has very different effects on *PMP2* and *DDR2*, primarily due to the effects of interactions.Presence of Haa1 reduces the regulatory effect of Skn7 while presence of Gis1 increases it. (B) Perturbation of Msn2 has very different effects on *HXK1* and *INO1*, as in panel A, however the independent effect of Msn2 on *HXK1* is much larger than that of Skn7 on *DDR2*. (C) Cyc8 interacts with multiple other TFs, including Gal 11 and Mig1, where it increases their effects on the target genes shown. The effects of Gal11 and Mig1 on *HXT6* are primarily due to an interaction with Cyc8, while Mig1 has a significant independent effect on *MAL33*.

## METHODS

### Transposon Calling Cards: Experiments

The calling cards assays were performed as previously described,^40^ with minor variations. Rather than expressing the Sir4-tagged TF from a plasmid, we C-terminal tagged the native TF-encoding gene using the CSWAT strain collection^41^ and deleted the native copy of *SIR4*. We also used barcoded plasmids containing the Tye5 transposon driven by the *GAL1-10* promotor and the *HIS3* and *URA3* auxotrophic markers (pRM1001-series as described in ref.^42^). An otherwise-empty plasmid carrying *LEU2* was used to confer leucine prototrophy.

### Transposon Calling Cards: Bioinformatics

Raw sequencing libraries are processed using the nf-core/callingcards pipeline (https://nf-co.re/callingcards/1.0.0/). Promoter enrichment is the normalized read count for a promotor in the experiment divided by the normalized read count in control samples lacking Sir4. We calculated it using the routine ‘yeast_promotor_enrichment’ from callingCardsTools (https://github.com/cmatKhan/callingCardsTools/tree/main). This also outputs a P-value for each promotor. Where there are multiple passing replicates of the same TF, the insertions from each replicate are summed and promoter enrichment is calculated on the summed replicate set.

### Calculation of Log Rank Binding and Log Rank Response signals

LRB and LRR are calculated as described in Results with an adjustment to spread values out over the full range when there are lots of ties (see Supplemental Methods). Briefly genes are ranked by binding *P*-value or response log-fold change, with the least significant genes given rank 1. For binding *P*-values, ties are broken by promotor enrichment. LRB and LRR are the logarithms of these ranks (base 10). This results in a distribution of LRB and LRR values that is approximately exponential. Genes whose promotors were present on plasmids during the Calling Cards procedure were removed from the dataset, as were those that did not respond to perturbation of any TF in both the ZEV data (used for modeling) and the TFKO data.^39^.

### Calculation of response rates

To calculate the response rate for each TF, we first identified the 25 genes that it binds most strongly. We consider a gene to be responsive to perturbation of a TF in the TF overexpression dataset (https://idea.research.calicolabs.com/data) if the field “log2_shrunken_timecourses” for the 15-minute time point is not zero.

### Interaction Feature Selection Pipeline

Separate models were fit for responses to perturbation of each TF using 1,000 random bootstrap samples of genes. For each sample, genes were divided into 4 folds by stratified random selection based on their LRB (Supplemental Methods). Using cross-validation, we trained models on 3 of the 4 folds and tested on the held out fold to assess R^2^. The optimal regularization parameters for the four folds were averaged, the model was refit on the whole dataset. The empirical distribution of values for each coefficient across all 1,000 bootstraps was used to test whether one-sided confidence intervals for the coefficient contained 0. For example, the 1-sided 95% confidence interval contains 0 if the bottom 5 percent of the empirical distribution includes zero or the top 5 percent of the empirical distribution includes zero.

## DISCUSSION

A fundamental question in regulatory systems biology is why some genes in whose promotors a TF binds respond to perturbations of that TF while others do not. One possible explanation was that ChIP experiments do not accurately report the binding locations of TFs. To investigate that possibility, we created a new, comprehensive dataset of TF binding locations in yeast using the Transposon Calling Cards (TCC) method. A higher fraction of genes whose promotors a TF binds responded to perturbations of the TF in our data, compared to older ChIP-based binding data. Nonetheless, many of the genes that TFs bound most strongly still did not respond to perturbations of the TF.

We hypothesized that functional synergy among bound TFs might explain why a TF’s binding in some promotors has a much greater effect on transcription than equally strong binding in other promotors. To investigate this, we fit mathematical models for predicting responses to TF perturbations from TF binding data. These models use the binding locations of all TFs, not just the perturbed TF (pTF), via interaction terms representing functional synergy. After rigorous statistical testing, we found many highly significant interactions between the binding signals from pTFs and the binding signals from other TFs, which we call modifier TFs (mTFs). Many different TFs served as modifiers, but the presence of Gis1, Opi1, Msn2, and Gal4 modified responses to the largest number of pTFs.

A few studies show experimental evidence of functional synergy among the activation domains of specific TF pairs. For example, activation domains from cMyc and SP1 were individually fused to the same artificial DNA binding domain and used to drive a fluorescent reporter from a synthetic promotor containing a single binding site for these synthetic TFs. When two plasmids expressing the two constructs were co-transfected into K562 cells at equal concentration, they drove expression from the artificial promoter at a higher level than when either plasmid was transfected at twice the concentration.^43^ Since there was only a single binding site, cooperative binding was ruled out. When a single fusion protein was made by linking two different activation domains to a single DNA binding domain, the effect on reporter gene expression was sometimes larger than the effects of two copies of the same domain.^44^ The reporter contained multiple binding sites so cooperative binding could explain this result, but only if there is heterotypic (but not homotypic) binding between the two activation domains. The mathematical modeling results we reported here complement and support this previous work and suggest that functional synergy may be more common than previously suspected.

Several possible explanations for our findings can be ruled out. One hypothesis is that the interaction terms in our models are serving as proxies for indirect effects of the pTF that are mediated by its effects on transcription of the mTF. If that were the case, the mTF would affect transcription of genes whose promotors it binds in regardless of whether the pTF is bound there. However, this is unlikely because the interaction terms were significant even when the main effects of the mTFs were included in the model. Our observations cannot be explained by effects of the mTF’s presence on the pTF’s binding because the pTF’s binding is measured, so any such effects are already accounted for in the univariate models using binding data on the pTF. They cannot be explained by the pTF being inactive in the experimental conditions used because this predicts no genes would respond to perturbations of the pTF; in our data, some genes responded to every perturbation. An explanation for the non-responsiveness of genes to deletion of a TF bound in their promotors is that the pTF is “redundant”. In other words, deleting it makes room for another TF that binds the same sites and has the same effects on transcription.^12^ However, that does not apply when the TF is perturbed by overexpression rather than gene deletion. An explanation that could apply to over-expression is that it does not affect promotors with the strongest binding signal from the pTF because all binding sites for the pTF are already occupied. This would predict that the strongest response would come from promotors with a moderate binding signal. However, we do not see this in our data.

There are several mechanisms by which functional synergy might work. The simplest is a threshold on regulatory potential – that is, on the total activating or repressing strength of all bound TFs. If strong binding of the pTF does not provide sufficient regulatory potential to pass the threshold for a significant effect on transcription, the presence of an mTF might be enough to push past the threshold. In one version, it does not matter which other TFs are present, so long as they have the same valence as the pTF – activating mTFs for an activating pTF and repressing mTFs for a repressing pTF. However, our data suggest that the pTF-mTF relationships are at least somewhat specific. In another version, the pTF and mTF cooperatively recruit some third factor, whose binding has not been measured and is therefore not accounted for in the univariate models. For example, several TFs including Mig1 and Nrg1 can recruit Cyc8 to promotors where they are bound. If Cyc8 binding were not measured, this could lead to a significant interaction between Mig1 and Nrg1. Since we measured Cyc8 binding, we see instead an interaction between Mig1 and Cyc8. Another possible mechanism is that the pTF and mTF might regulate different phases of a transcriptional cycle, such that both are needed for productive transcription.^43,45^ Another is that overexpression of a pTF might have a greater effect on promotors where it can displace an mTF that is bound with high affinity at sites that the pTF binds with lower affinity. More research is needed to determine the mechanism of functional synergy for each pTF-mTF pair.

A longstanding goal in regulatory genomics has been to develop algorithms that can map TF networks using gene expression data, binding data, or DNA sequence motifs.

Recently, several of these information sources have been combined.^46,47^ These maps are typically represented as a list of relationships (“edges”) between TFs and the genes they regulate, each with an associated score representing the algorithm’s confidence that the TF in fact regulates the target. While many such algorithms have been published over more than two decades, there are fundamental limitations to this conception of transcriptional regulation. First, as we have shown, TF-target relationships are not binary – they can be modified by interactions between TFs. Second, TF network maps typically represent the potential of a TF to regulate a target gene, rather than asserting that the TF is actively modifying the expression of the gene in given biological condition, cell type, or sample.

The models we have fit, on the other hand, are functions that map the binding signals of TFs, measured in a specific biological condition, to the predicted effects of the TFs on the expression of target genes. These predicted effects can naturally be thought of as edge scores in a condition-specific network map. The predicted effect of a modifier TF on the response can be depicted as an activating or inhibiting effect on the primary edge (Fig. 4). These maps are based on both binding data and expression data, since both were used to fit the models. Unlike maps built from perturbation data alone, they will have high-scoring edges from a TF to a gene only if the TF binds in the gene’s promotor. New maps for new growth conditions can be constructed by applying the trained models to binding data from those conditions. However, those models will not account for changes in TF activity or changes in synergistic partners. If both binding and perturbation response data for the new condition can be obtained, new models can be fit to the data before extracting networks.

Computational models of transcriptional regulation can be classified by their inputs (predictors) and outputs (response variables). Much effort has been devoted to sequence-to-binding models and sequence-to-expression models, yielding substantial advances in predictive power. Comparatively little work has been done on models that use measured TF binding locations as input, and almost none on models that use binding data to predict expression changes in response to TF perturbations. Recently, neural networks (NNs) have become a popular formalism for sequence-to-binding and sequence-to-expression models. This is feasible because very large datasets exist for these tasks. For example massively parallel reporter assays (MPRAs) measure gene expression driven by synthesized artificial promotors, millions of which can be manufactured. However, no datasets exist in which binding locations and perturbation responses have been measured for large numbers of synthetic promotors. Yeast has only ∼6,000 genes, yielding ∼6,000 responses to perturbation of each TF. This small training sample limits the applicability of deep learning to this problem. Until adequate training data are available, the generalized additive models framework will be the method of choice for predicting which TF binding sites are functional and which are not. Within that framework, future work can explore more expressive models. One example is models with terms for the indirect effects of TF perturbation. Another is models that include non-linear effects of interaction terms, analogous to the cubic model we used for the independent effect perturbed TF binding on genes’ responses. Such efforts may bring us even closer to the goal of accurately predicting how the genes whose promotors a TF binds will respond to perturbations of that TF.

## Supporting information

Log Rank Binding signal

Log Rank Response signal

Coefficients from cubic models

Residuals from cubic models

Interaction terms surviving Stage 1

: Interaction terms surviving Stage 2

Interaction terms surviving Stage 3

