## Supplementary material for "UBIQUITOUS FUNCTIONAL SYNERGY PARTIALLY EXPLAINS WHY MOST TRANSCRIPTION FACTOR BINDING IS NON-FUNCTIONAL": Online supplement

SUPPLEMENTAL FILES

- File S1: Log Rank Binding signal (LRB) for all samples used in the modeling.
- File S2: Log Rank Response signal (LRR) for all samples used in the modeling.
- File S3: Coefficients (including intercept) from cubic models.
- File S4: Residuals from cubic models.
- File S5: Interaction terms surviving Stage 1, with confidence intervals on coefficients, median coefficient across bootstraps, and coefficient when all terms surviving Stage 1 are fit to the full, original dataset by LASSO.
- File S6: Interaction terms surviving Stage 2, with confidence intervals on coefficients, and median coefficient across bootstraps.
- File S7: Interaction terms surviving Stage 3.

SUPPLEMENTAL METHODS

Transposon Calling Cards: Bioinformatics

Raw sequencing libraries are processed using the nf-core/callingcards pipeline (https://nf-co.re/callingcards/1.0.0/) with the setting -profile default_yeast and all other settings left at default values. The `default_yeast` profile uses bwamem2 version 2.2.1 and aligns to the *s. cerevisiae* sacCer3 genome. Promoter enrichment was calculated on the output of the nf-core/callingcards workflow using annotations from SGD version R64-3-1_20210421 and the routine `yeast_promotor_enrichment`, part of the command line interface of callingCardsTools (<https://github.com/cmatKhan/callingCardsTools/tree/main>). One of the outputs is a *P*-value for each promotor, based on the null hypothesis that the rate of transposon insertion in that promotor is no greater than in a control dataset from a strain lacking Sir4. Promotors are ranked by their *P*-value and compared to perturbation response datasets using Dual Threshold Optimization (DTO; {Kang, 2020, 32060051}; https://github.com/BrentLab/Dual_Threshold_Optimization). DTO outputs an empirical *P*-value for the null hypothesis that there is no relationship between the genes that are highly ranked in the binding data and those that are highly ranked in the perturbation response data. Each Calling Cards replicate was considered QC-passing if it had at least 3,000 insertions and a DTO empirical *P*-value <= 0.01 when compared to perturbation response data from either ref.{Kemmeren, 2014, 24766815} or ref.{Hackett, 2020, 32181581}

Where there are multiple passing replicates of the same transcription factor, the insertions are merged and promoter enrichment is calculated on the merged replicate set. The promoter regions over which insertions are quantified may be found at <https://huggingface.co/datasets/BrentLab/yeast_genome_resources/blob/main/brentlab_promoters.bed>. Processed data (qbeds, describing calling cards insertion locations) are available at (GEO).

Calculation of Log Ranking Binding (LRB) signal

LRB can be calculated for any binding profile that includes a *P*-value for each gene and an enrichment score for each gene reflecting the enrichment of reads over the gene’s promotor, relative to a control experiment. Such profiles can be produced from many sequencing-based assays including Calling Cards, ChIP-seq, ChIP-exo, or ChEC-seq. To calculate LRB from the binding profile of a TF, genes are ranked by *P*-value, with the least significant *P*-value given rank one and ties broken by enrichment. If genes have the same *P-*value and the same enrichment they are given the same rank. After ranking, we took the logarithm of the rank. The resulting distribution of LRB values is approximately exponential.

Genes with identical P-value (typically *P*=0 or *P*=1) and identical enrichment (mostly 0 counts for genes with *P* = 1) remained tied. When ranking, all elements within a tied group receive the same rank. Typically, they would be assigned the minimum, maximum, or mean of the block of ranks occupied by the tie. However, when there are large numbers of tied elements at the top and bottom rankings, that approach can substantially compress the range of ranks used. To reduce this compression, we applied a formula that picks which of the ranks occupied by the tie to assign to the tied elements that depends on where the tied elements lie in the overall ranking.

$$assignedrank=minrank+ntied\frac{\mathrm{averagerank}}{\mathrm{ngenes}}$$

where $\mathrm{minrank}$ is the minimum rank of the block occupied by the tie, $\mathrm{averagerank}$, is the average of the block occupied by the tie, $\mathrm{ntied}$ is the number of genes in the tie, and $\mathrm{ngenes}$ is the total number of yeast genes. For example, if there are 20 genes and 5 of them are tied for ranks 1-5, they would receive the rank 1+5*(3/20) = 1.75, less than their average rank of 3. If there are 20 genes and 5 of them are tied for ranks 15-20, they would receive the rank 15+5*(17/20) = 19.25, more than their average rank of 17.

Calculation of the Log Rank Response (LRR)

LRR can be calculated for any perturbation response profile. It is calculated the same way as LRB except that genes are ranked by log fold change relative to the unperturbed control. There are very few ties in log fold changes, so no tie-breaking mechanism was needed.

Blacklisted target genes

After intersecting binding and perturbation response target gene features, we excluded five genes that are either engineered in the calling cards base strain or located at divergent promoters with engineered genes (*URA3*/YEL021W, *HIS3*/YOR202W, *MRM1*/YOR201C, *LEU2*/YCL018W, and YOR203W). We also excluded 80 genes that did not respond to perturbation of any TF in both the ZEV data (used for modeling) and the TFKO data.{Kemmeren, 2014, 24766815} Supplemental Files S1 and S2 do not contain entries for these genes.

Public Datasets

The Harbison binding data were downloaded from <http://younglab.wi.mit.edu/regulatory_code/GWLD.html>. The pvalue and ratio data were downloaded, and both TF and target annotations were harmonized with SGD S288C_R64-3-1 (see <https://huggingface.co/datasets/BrentLab/harbison_2004/blob/main/scripts/parse_harbison_data.R> for details).

The TFKO data {Kemmeren, 2014, 24766815} was downloaded from <https://deleteome.holstegelab.nl/>. The TF mutant and target feature names were harmonized with the SGD S288C_R64-3-1 annotation file (see https://huggingface.co/datasets/BrentLab/kemmeren_2014/blob/main/scripts/parse_kemmeren_data.R for details). For the perturbation effect, we use the data with the slow growth effect computationally removed (also provided by the author at their website). For calculating response rates (Fig. 1E) we label a gene as responsive if the p-value is <= 0.05. Where more than one probe is assigned to the same feature, we use assign the feature the largest effect (Madj) or the smallest p-value. The TF overexpression data {Hackett, 2020, 32181581} was downloaded from <https://idea.research.calicolabs.com/data> and harmonized to SGD S288C_R64-3-1. We used the data from time point 15. For regression, we used the field “log2_ratio” as the effect. For calculating response rates, we label a gene as responsive if the field “log2_shrunken_timecourses” is not zero. The script used to parse the data, and the data itself can be found at <https://huggingface.co/datasets/BrentLab/hackett_2020>.

Univariate linear models

To fit linear models without data transformation, we calculated the binding signal strength as the log 2 fold change (LFC) of transposon insertion frequency in each genes promoter compared to the insertion frequence in the same promoter in [specify control experiment experiments]. Promoters defined to be [promoter definition]. We also calculated the non-shrunken LFC of each gene 15 minutes after inducing overexpression of each TF by estradiol addition in engineered ZEV strains [cite ZEV paper] (see Public Datasets). We fit univariate linear regression models to predict the perturbation response from the binding signal by least squares.

Interaction Feature Selection Pipeline

Separate models were fit for responses to perturbation of each TF using 1,000 random bootstrap samples of genes. Genes were divided into 4 folds by stratified random selection based on their LRB. Three strata were defined by LRB ranks (1, 64], (64, 512], and (512, max], where 1 represents the strongest binding signal. Using cross-validation, we trained models on 3 of the 4 folds and tested on the held out fold to assess R^2^. For each fold, the optimal value of the regularization parameter lambda was chosen by using the StratifiedKFold function from scikit-learn and the resulting 4 lambdas were averaged. Fixing lambda at the average, the model was refit on the whole datasets and the coefficients for each term (including 0 coefficients) were returned. This produced an empirical distribution of coefficient values for each coefficient across all 1,000 bootstraps. These were used to test whether one-sided confidence intervals for the coefficient contained 0. For example, the 1-sided 95% confidence interval contains 0 if the bottom 5 percent of the empirical distribution includes zero or the top 5 percent of the empirical distribution includes zero.

Stage 3 of the pipeline used LASSO to fit a model containing interaction terms that survived Stage 2 along with the corresponding independent (main) effect terms. Any interaction term whose coefficient went to zero or changed sign in the presence of the corresponding independent effect was dropped.

SUPPLEMENTAL FIGURES

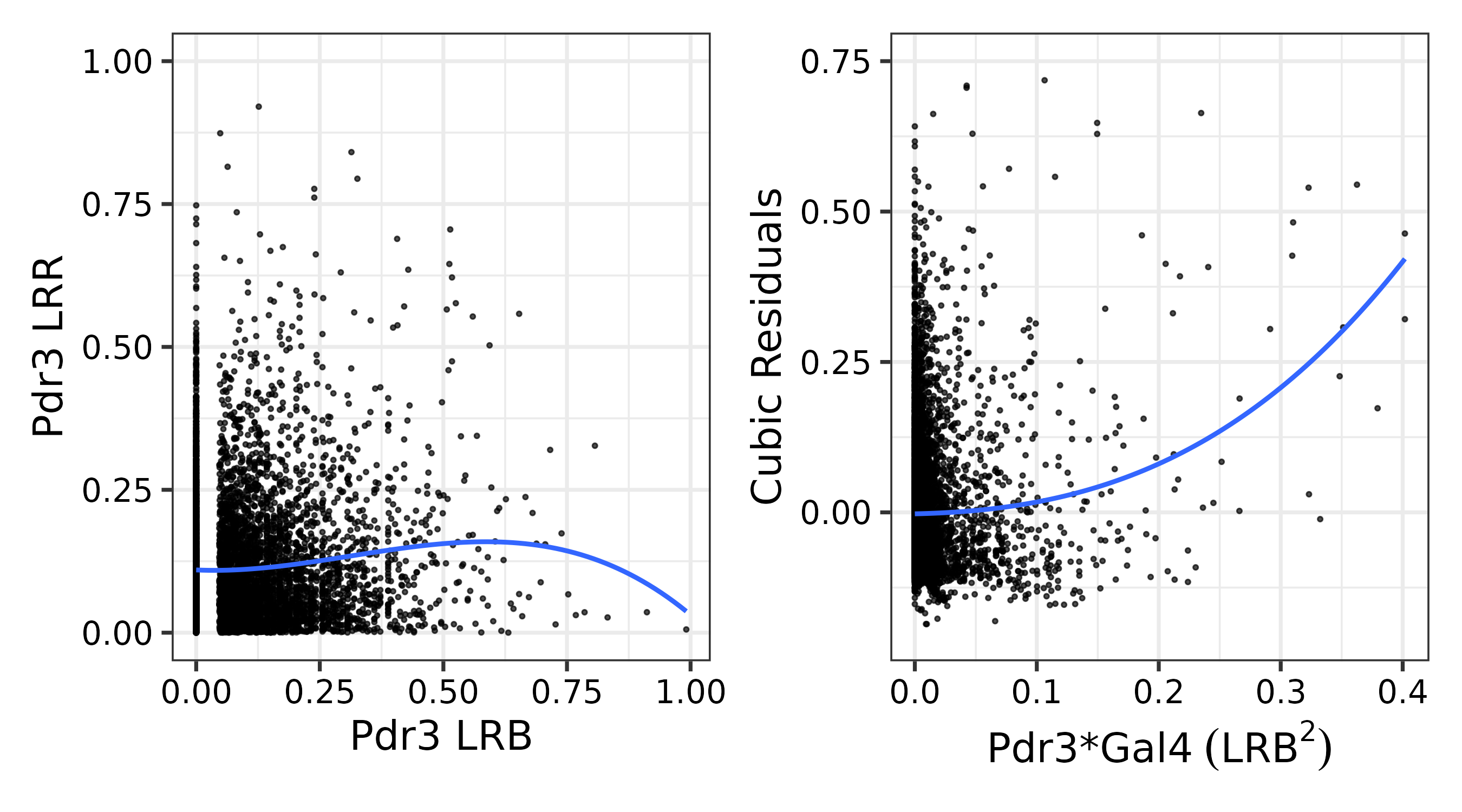

**Figure S1** Left: The cubic polynomial fit to the LRR of all genes in response to perturbation of *PDR3* (Y-axis) as function of Pdr3 binding strength (X-axis). This fit explains almost none of the variance in the data (a linear fit would explain even less). The distance between each point and the line is the residual error, which cannot be explained by Pdr3 binding strength. Right: The residual for each gene as a function of the product of Pdr3 and Gal4 binding strengths. The fact that this line is not horizontal shows that this product explains some of the residual variance that cannot be explained by the independent effect of Pdr3 binding.
